# Adaptive introgression: an untapped evolutionary mechanism for crop adaptation

**DOI:** 10.1101/379966

**Authors:** Concetta Burgarella, Adeline Barnaud, Ndjido Ardo Kane, Frédérique Jankowsky, Nora Scarcelli, Claire Billot, Yves Vigouroux, Cécile Berthouly-Salazar

**Affiliations:** IRD, UMR DIADE, Montpellier, France; Université de Montpellier, Montpellier, France; CIRAD, UMR AGAP, F-34398 Montpellier, France; AGAP, Univ Montpellier, CIRAD, INRA, Montpellier SupAgro, Montpellier, France; Institut Sénégalais de Recherches Agricoles (ISRA), LNRPV, Dakar, Senegal; Laboratoire Mixte International LAPSE, Dakar, Senegal; CIRAD, UPR GREEN, F-34398, Montpellier, France; GREEN, CIRAD, Univ Montpellier, Montpellier, France; Institut Sénégalais de Recherches Agricoles (ISRA), BAME, Dakar, Senegal

**Keywords:** crops, wild relatives, domestication, selection, gene flow, adaptive introgression, farmer’s practices

## Abstract

Global environmental changes strongly impact wild and domesticated species biology and their associated ecosystem services. For crops, global warming has led to significant changes in terms of phenology and/or yield. To respond to the agricultural challenges of this century, there is a strong need for harnessing the genetic variability of crops and adapting them to new conditions. Gene flow, from either the same species or a different species, may be an immediate primary source to widen genetic diversity and adaptions to various environments. When the incorporation of a foreign variant leads to an increase of the fitness of the recipient pool, it is referred to as “adaptive introgression”. Crop species are excellent case studies of this phenomenon since their genetic variability has been considerably reduced over space and time but most of them continue exchanging genetic material with their wild relatives. In this paper, we review studies of adaptive introgression, presenting methodological approaches and challenges to detecting it. We pay particular attention to the potential of this evolutionary mechanism for the adaptation of crops. Furthermore, we discuss the importance of farmers’ knowledge and practices in shaping wild-to-crop gene flow. Finally, we argue that screening the wild introgression already existing in the cultivated gene pool may be an effective strategy for uncovering wild diversity relevant for crop adaptation to current environmental changes and for informing new breeding directions.

## 1. Introduction

The fate of wild and domesticated species and their associated ecosystem services is increasingly depending on global environmental changes, as climate warming, nitrogen cycle alteration or land use (Perring et al., 2015; Shibata et al., 2015; Walther et al., 2002). For instance, modifications of temperature and rainfall regimes have been shown to directly impact on plant phenology or distribution within these past decades. A meta-analysis including nearly 1600 species showed that 41% of them had experienced phenological advancement and northward range movement (Parmesan and Yohe, 2003). In a mountain area, Crimmins et al. (2011) documented downward elevation shifts driven by water deficits in 64 plant species. Forest inventories across the eastern USA revealed direct effects of climate on forest biomass, through changes in tree species composition towards species more drought-tolerant, but slower growing. Climate effects have also been reported on major crop species. Global maize and wheat yield has declined by 4-6% since the early 1980s (Lobell et al., 2011). Earlier flowering and changes in genetic composition have been recorded in the staple African cereal pearl millet (Vigouroux et al., 2011). Reduced flowering time and loss of genetic diversity in response to increasing temperatures have also been observed in wild wheat and barley over less than 30 years (Nevo, 2012). Future climate scenarios foresee an acceleration of the rise in temperature and an increase in hydrological variability (IPCC, 2014), which are probably the prelude to further dramatic consequences for species biology.

Phenotypic plasticity and dispersal (through seeds and pollen) can be very rapid responses to change, but it is less clear whether adaptation, the evolution of genetic traits making organisms better fitted to survive and reproduce in their environment, could also play a significant role in this process. The ability to adapt to new conditions depends on the rate of environmental change (Loarie et al., 2009) and on the genetic variability available (Doi et al., 2010; Hoffmann et al., 2017). The genetic diversity of a population relies either on standing genetic variation or on new genetic variants. On short time scales, the mutation rate may be too low. Moreover, standing genetic variation may be limited, especially for populations whose genetic diversity has been reduced by demographic bottlenecks, as many domesticated species (Glémin and Bataillon, 2009). However, new variants could arise from gene flow, in numbers up to two or three orders of magnitude more than that introduced by mutation (Grant and Grant, 1994). Thus, gene flow, either from the same species or a different species, may amount to an immediate primary source of functional alleles (Ellstrand, 2014). If a foreign functional variant increases the fitness of the recipient pool, it increases in frequency across generations, a phenomenon referred to as “adaptive introgression” (Anderson, 1949; Rieseberg and Wendel, 1993).

Prior to any possible “adaptive introgression”, hybridization/gene flow events need to take place. However, the fate of hybrids is uncertain. Hybrids are usually selected against in parental habitats (Baack et al., 2015) because they show reduced fertility and viability (Lowry et al., 2008). This process might be due to genetic incompatibilities and/or the break-up of epistatic co-adapted gene complexes (Barton and Hewitt, 1985; Hewitt, 1988). Yet, when species are not too divergent and isolating barriers are incomplete (Coyne and Orr, 2004), hybridization can lead to the introgression of advantageous alleles (Barton and Bengtsson, 1986). Compared to neutral introgression, which could be lost by drift across generations, adaptive introgression would be maintained and may eventually give rise to fixation (Figure 1). Interestingly, hybridization has the potential to introduce large sets of new alleles simultaneously at multiple unlinked loci, allowing adaptation even for polygenic traits (Abbott et al., 2013; Mallet, 2007), thus playing a key role in species evolution (e.g. Arnold and Kunte, 2017; Arnold and Martin, 2009; Hedrick, 2013).

**Figure 1.**
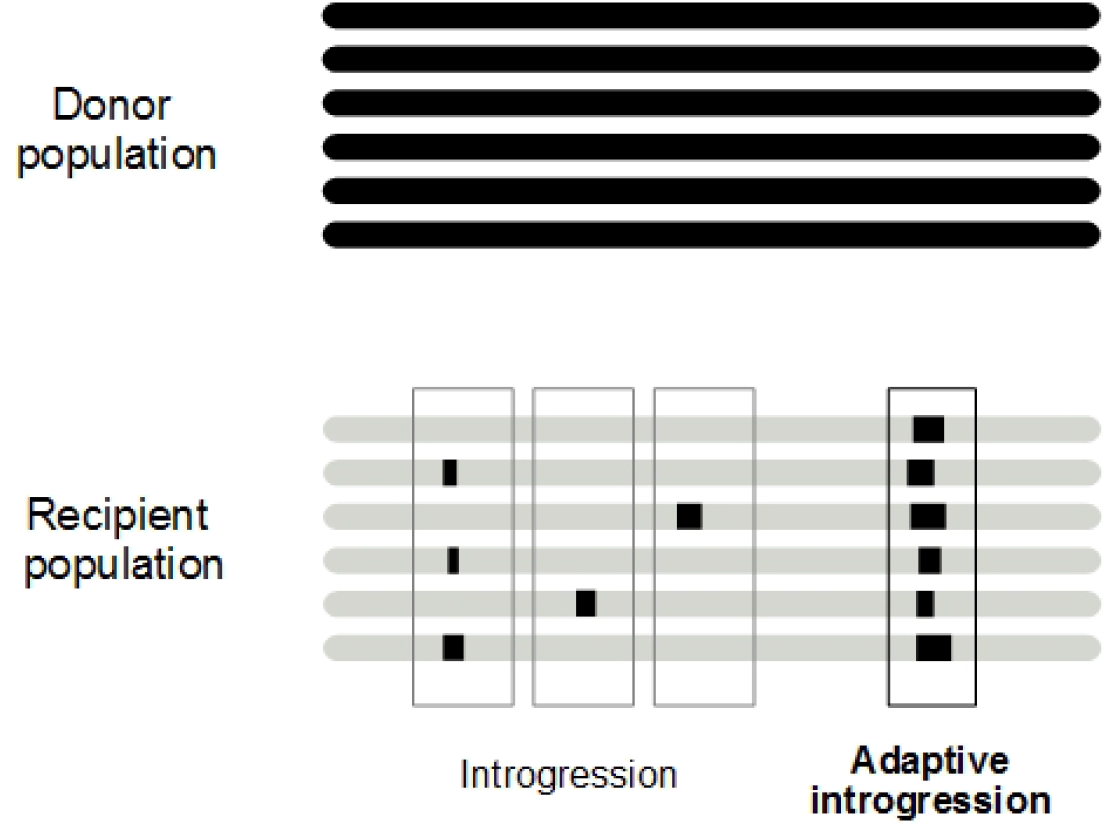
Genomic signature of adaptive introgression. A genomic fragment is transferred from the donor to the recipient population via gene flow and subsequent backcrosses. If it enhances the fitness of the recipient population, it experiences positive selection and rises in frequency.

In the context of species conservation and management a large body of literature discuss how gene flow could also be associated with negative effects. Undesirable consequences of gene flow are associated with the risks of invasiveness (Ellstrand and Schierenbeck, 2000; Whitney et al., 2006), transgene escape (Ellstrand et al., 2013 and references therein), or genetic erosion of native populations (Wolf et al., 2001). Little attention has been paid to the potential of managed gene flow to increase genetic variation for species rescue (Hedrick, 2009) and adaptation (Aitken and Whitlock, 2013). Up to now, the potential of adaptive introgression as a source of adaptation to on-going global changes for domesticated species has been overlooked.

In this paper, we review studies of adaptive introgression, paying particular attention to the potential of this evolutionary mechanism for the adaptation of crops. By the term ‘introgression’ we mean the consequence of gene flow, at both the interspecific and the intraspecific level, i.e. independently of the taxonomic classification of the gene pools exchanging genetic material. We focus on sexually reproductive organisms; and we did not directly discuss horizontal gene transfer (reviewed in Arnold and Kunte, 2017). Within this framework, we report on 34 case studies addressing adaptive introgression (Table 1). After discussing methodological approaches and challenges to detecting adaptive introgression, we focus on gene flow in crops and the importance of farmer practices in shaping wild-to-crop gene flow.

**Table 1.**
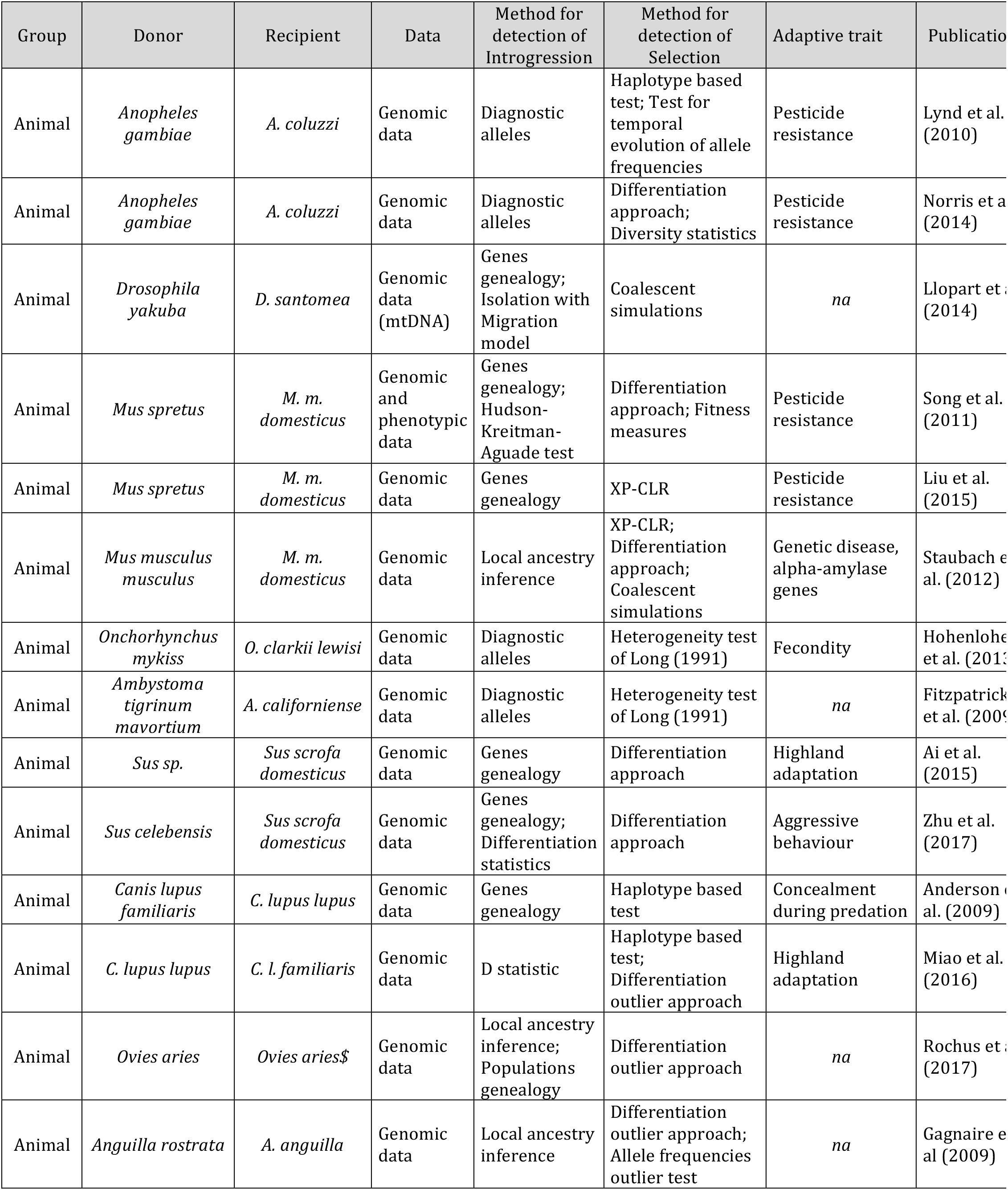
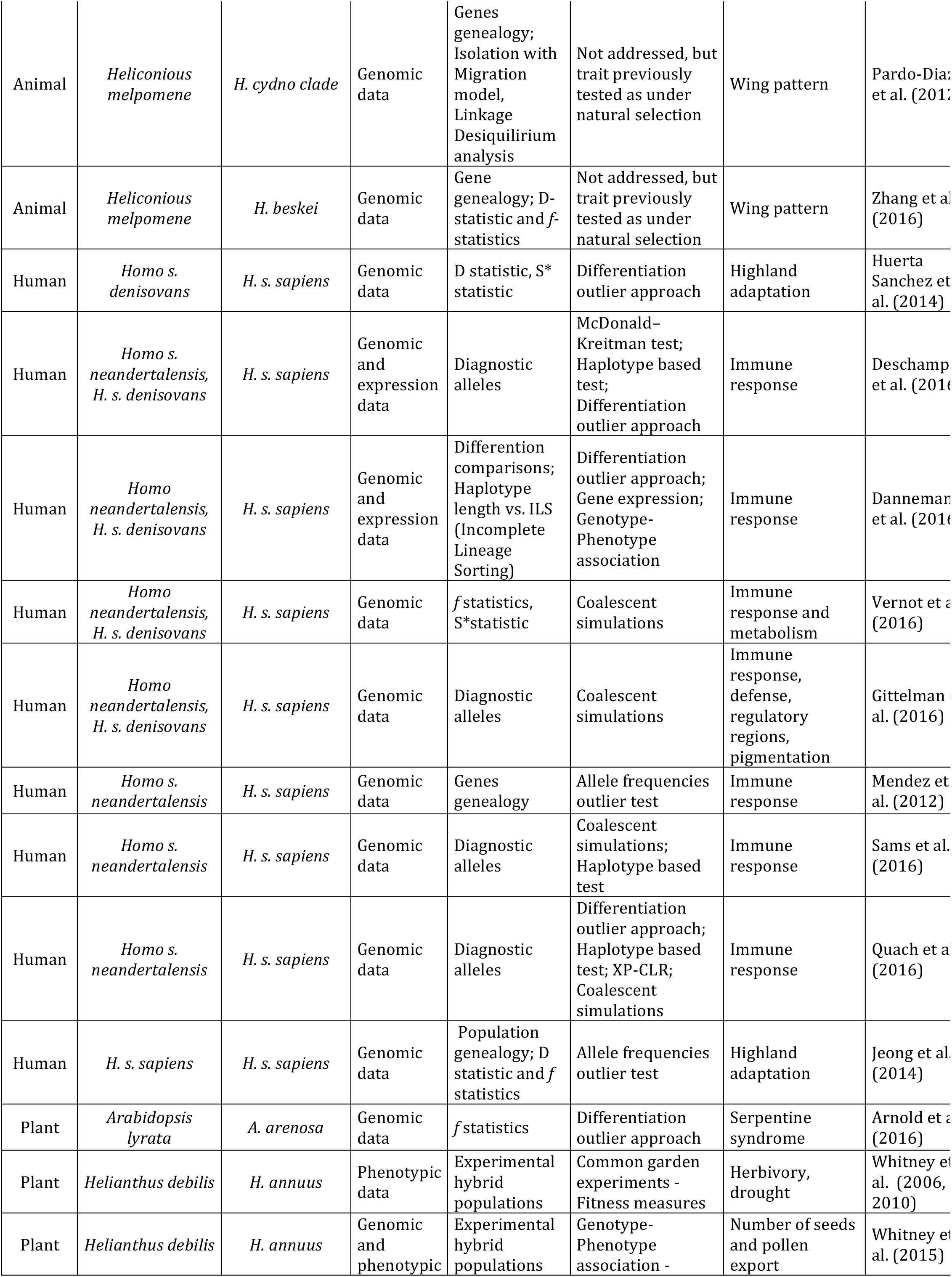
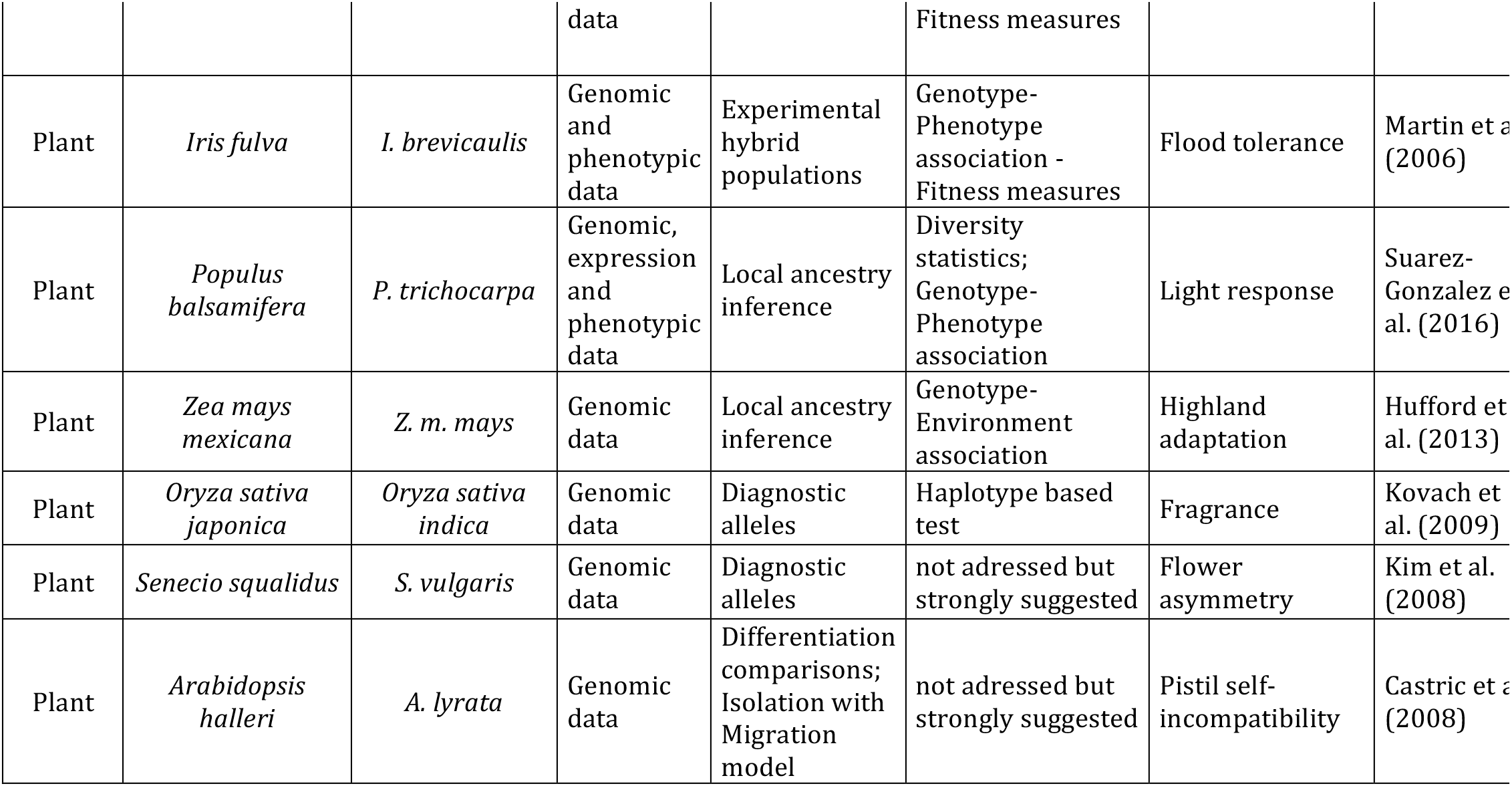
Summary of studies reviewed. Species names for donor and recipient taxa are listed, as well as the type of data and methods used for 1) detection of the introgression and 2) detection of the selection. “Genomic data” term include s whole-genome sequences or candidates genes sequencing. “Genetic data refers to molecular markers such as QTL or SSR.

## 2. Empirical evidence of adaptive introgression

Despite the occurrence of hybridization in nature, involving 25% of plants and 11% of animals (Mallet, 2005), relatively little evidence supports the fact that gene flow leads to enhanced fitness, i.e. adaptive introgression. This may be because investigating the fitness of adaptive introgression is intrinsically difficult. For example, demonstrations of adaptive introgression in *Helianthus* were based on crossing and backcrossing to experimentally reproduce introgression and demonstrate enhanced fitness in the recipient taxon (Whitney et al., 2006, 2010). With high-throughput sequencing technologies, we can instead search for selection signatures on the introgressed variant in the recipient genomes (e.g. Arnold et al., 2016; Racimo et al., 2015). An increasing number of publications involving large-scale genetic data are accumulating in this field (Table 1). Those studies reveal or confirm instances of adaptive introgression in many kinds of organisms, including domesticated species (e.g. Anderson et al., 2009; Hufford et al., 2013; Kovach et al., 2009; Miao et al., 2016; Rochus et al., 2017). It also appears that different abiotic and biotic selective pressures drive the introgression of adaptive traits, with evolutionary consequences spanning different spatial and temporal scales.

Recent striking studies report ancient adaptive introgression events in animals or humans. Up to 5% of the modern human genome might be of introgressed origin from archaic hominins (e.g. Green et al., 2010; Hsieh et al., 2016; Reich et al., 2009; Sankararaman et al., 2014). Introgressed loci from Neanderthals or Denisovans appear to be linked to skin colour phenotypes (Vernot and Akey, 2014), immune responses (Abi-Rached et al., 2011) and hypoxia adaptation to high altitudes (Huerta-Sánchez et al., 2014). In animals, Tibetan Mastiff dogs were also found to have received a genomic region encompassing two genes linked to hypoxia adaptation (the *EPAS1* and the *HBB* genes) from the local populations of grey wolf (Miao et al., 2016; Zhang et al., 2014). On the other hand, the American grey wolf inherited an adaptive allele for coat pigmentation from past hybridization with domestic dogs (Anderson et al., 2009). In Chinese and European pigs, gene flow would seem to date from the Pleistocene era, when northern Chinese and European breeds would appear to have acquired a large adaptive genomic region from a *Sus* species that is now extinct (Ai et al., 2015).

Adaptive introgression can also take place on a much shorter evolutionary time scale. For instance, the acquisition of pesticide resistance through genetic exchange can be achieved in a few decades, as observed in insects (Lynd et al., 2010; Weetman et al., 2010) and rodents (Liu et al., 2015; Song et al., 2011). A mutation in the voltage-gated sodium channel gene (*kdr*) provides strong resistance to pyrethroid and dichlorodiphenyltrichloroethane (DDT) in the mosquito *Anopheles gambiae*, one of the vectors of malaria in sub-Saharan Africa. The resistance allele was transferred from *A. gambiae sensu stricto* (S form) to a conspecific *A. coluzzi* (M form) (Weill et al., 2000) and its frequency has greatly increased in *A. coluzzi* populations over the past two decades (Lynd et al., 2010; Norris et al., 2015). Selection tests and demographic simulations suggest this increase has been driven by selection (Lynd et al., 2010; Weetman et al., 2010). Another recent example of adaptive introgression in plants has been discovered in a population of *Arabidopsis arenosa* (Arnold et al., 2016) using genome scans. Several genomic signatures of selection associated with adaptation to serpentine soils were found to be of introgressed origin from a different species, *A. lyrata*. The uncontrolled escape of agricultural adaptations (e.g. resistance to biotic and abiotic stressors, often achieved with transgenes) from fields to wild populations, through weedy forms, is another well-known case of rapid adaptive introgression. This causes substantial yield losses and requires strong economic efforts in managing cultivated and wild species (e.g. Ellstrand et al., 2013; Hooftman et al., 2007; Rose et al., 2009; Uwimana et al., 2012).

## 3. Characterizing adaptive introgression with genetic data

To infer adaptive introgression, it is necessary to demonstrate 1) the introgression, by showing the foreign origin of the genetic variant and its persistence in the recipient pool (i.e. should be found in backcrossed generations), and 2) its adaptive value, by identifying selection footprints on the introgressed fragment and (if possible) its fitness value. Genomic studies of adaptive introgression seek to aim at gathering these three lines of evidence. Because of multiples factors (migration rate, number of generations since introgression, intensity of selection) affecting the introgression process and its interaction with selection, a variety of genomic patterns can be observed in the recipient population (Figure 1). As these are complex patterns, there is no unique approach to detecting signatures of adaptive introgression (Table 1). Below, we detail some of the most common approaches used to detect introgression and to prove the action of selection (cf. Table 1).

### 3.1 Detection of introgression

The aim of detecting introgression is to identify populations and individuals of admixed origin and quantify rates of gene flow, but also to find the traits or the genomic regions that have crossed isolation barriers. Availability of whole genome data maximizes the chances of detecting introgression even when it is rare in the genome (Hufford et al., 2013; Racimo et al., 2015; Rochus et al., 2017; Schaefer et al., 2016). In the following sections, we describe approaches used to detect introgression with genetic data, bearing in mind that none of them provides absolute proof of introgression and that the best strategy is to gather evidence in different ways.

The ability to detect introgression increases with the divergence between the hybridizing taxa. For higher divergent taxa, we have more markers fixed between species or with large allele frequency differences. These “diagnostic alleles” allow easy identification of the ancestry of a genomic fragment in the recipient population (e.g. Gittelman et al., 2016; Kim et al., 2008; Kovach et al., 2009; Norris et al., 2015; Smith et al., 2004). However, even with slight differences in allele frequencies, genetic clustering methods can be used to identify introgression. A variety of approaches are available, such as multivariate analyses (e.g. Frichot et al., 2014; Jombart et al., 2009) or Bayesian algorithms (e.g. Alexander et al., 2009; Anderson and Thompson, 2002; Pritchard et al., 2000) (Figure 2a). The power of these global ancestry methods to detect gene flow comes from the use of multiple independent (i.e. not physically linked) polymorphic markers. These methods can be applied both genome-wide (Gagnaire et al., 2009; e.g. Rochus et al., 2017) and to single genomic regions. Window-based analyses of ancestry along the maize genome were successfully used to identify introgressed fragments from the wild progenitor teosinte (Hufford et al., 2013).

**Figure 2.**
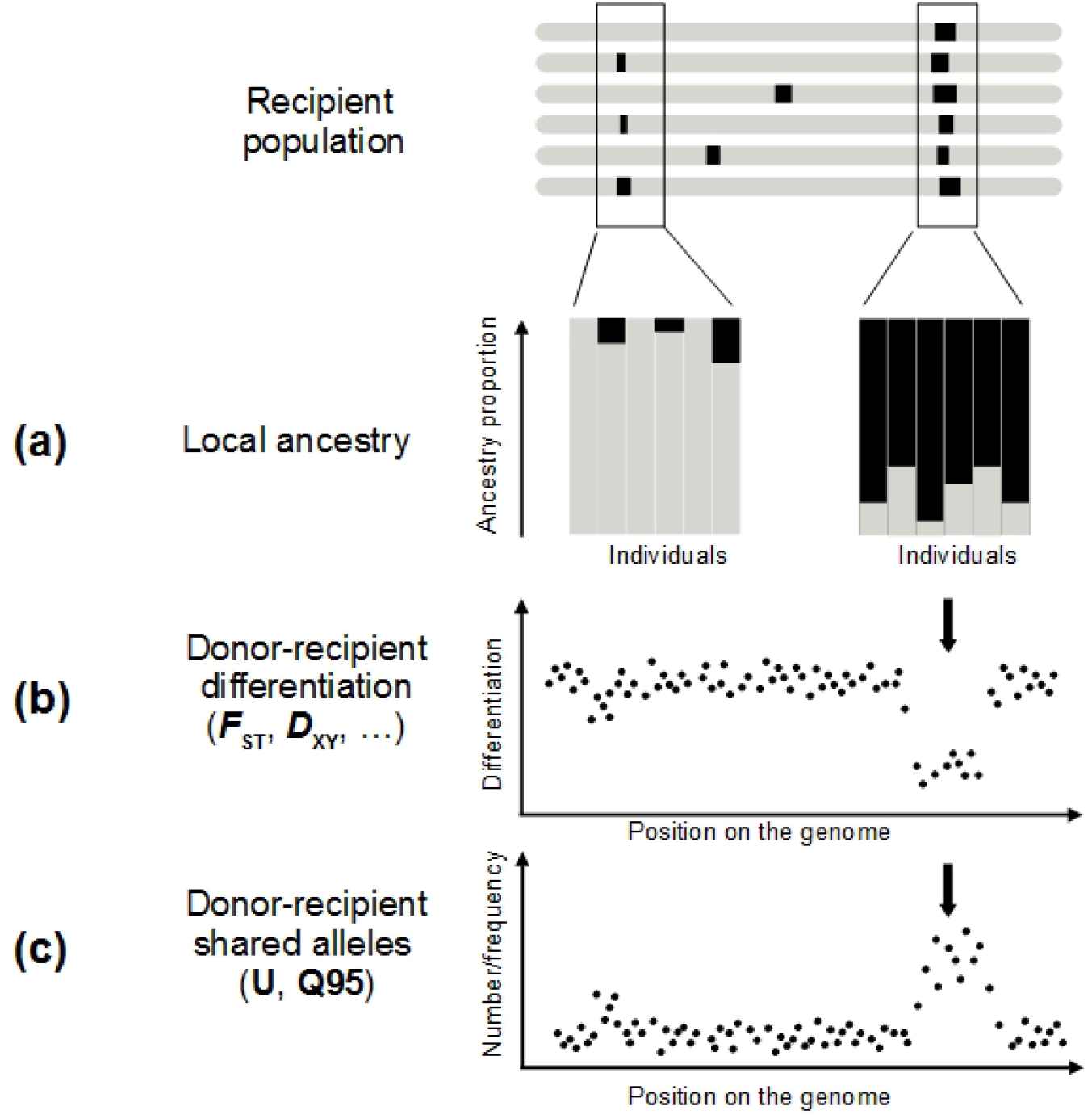
Approaches to detect introgression (a, b) and adaptive introgression (c). Regions of donor origin in the recipient genome can be revealed by performing local ancestry analyses (a) and comparisons of donor-recipient differentiation levels (b). Introgressed fragments will show a larger proportion of ancestry in the donor population (a. in black) and lower differentiation (b. arrow) than non-introgressed recipient genomic regions. Positive selection increases the frequency of the donor allele and the neutral variants physically linked to it. The result is a local higher number and frequency of alleles shared by donor and recipient populations and absent in other non-introgressed populations (c. arrow).

When higher density molecular markers are available, other recent methods are able to assign an ancestry probability to each polymorphic variant (Racimo et al., 2015; Schaefer et al., 2016). These local ancestry methods use probabilistic approaches, such as Hidden Markov Models (e.g. Reich et al., 2012), or Conditional Random Fields (e.g. Sankararaman et al., 2014) to infer the ancestry state of each site, taking into account the information of physically close positions. As physical linkage disequilibrium patterns dilute with generations, these approaches are less efficient for the detection of ancient introgression, compared to global ancestry methods. While some implementations require phased data (e.g. Song and Hein, 2005) or training data (e.g. Sankararaman et al., 2014), more recent developments have overcome these constraints (e.g. Guan, 2014). So far, such approaches have been mainly applied to model species (e.g. Staubach et al., 2012 on *Mus musculus*; Turissini and Matute, 2017 on *Drosophila*; Zhou et al., 2016 on humans), but the increasing availability of whole genome data will soon make them suitable for other study systems.

The approaches described above help to quantify the amount of shared diversity between genetic pools. Shared variants between populations may be the result of different processes other than introgression: the retention of ancestral polymorphic alleles by chance (referred to as Incomplete Lineage Sorting, ILS, Figure 3), balancing selection or convergence (see Hedrick, 2013 for a comparison). For lower divergence times (as for wild-crop complexes), the probability that the two related groups have conserved ancestral polymorphism is higher. Thus, in most cases, the main challenge to detecting introgression is to distinguish it from ancestral shared polymorphism. Tracking the absence of the introgressed variants in ancient samples of the recipient pool would be an efficient way of excluding shared ancestral polymorphism. However, historical samples are difficult to obtain for most biological systems, so different methods have been developed to search for specific signatures on the genome that help to differentiate between introgressed fragments and inherited ancestral fragments.

**Figure 3.**
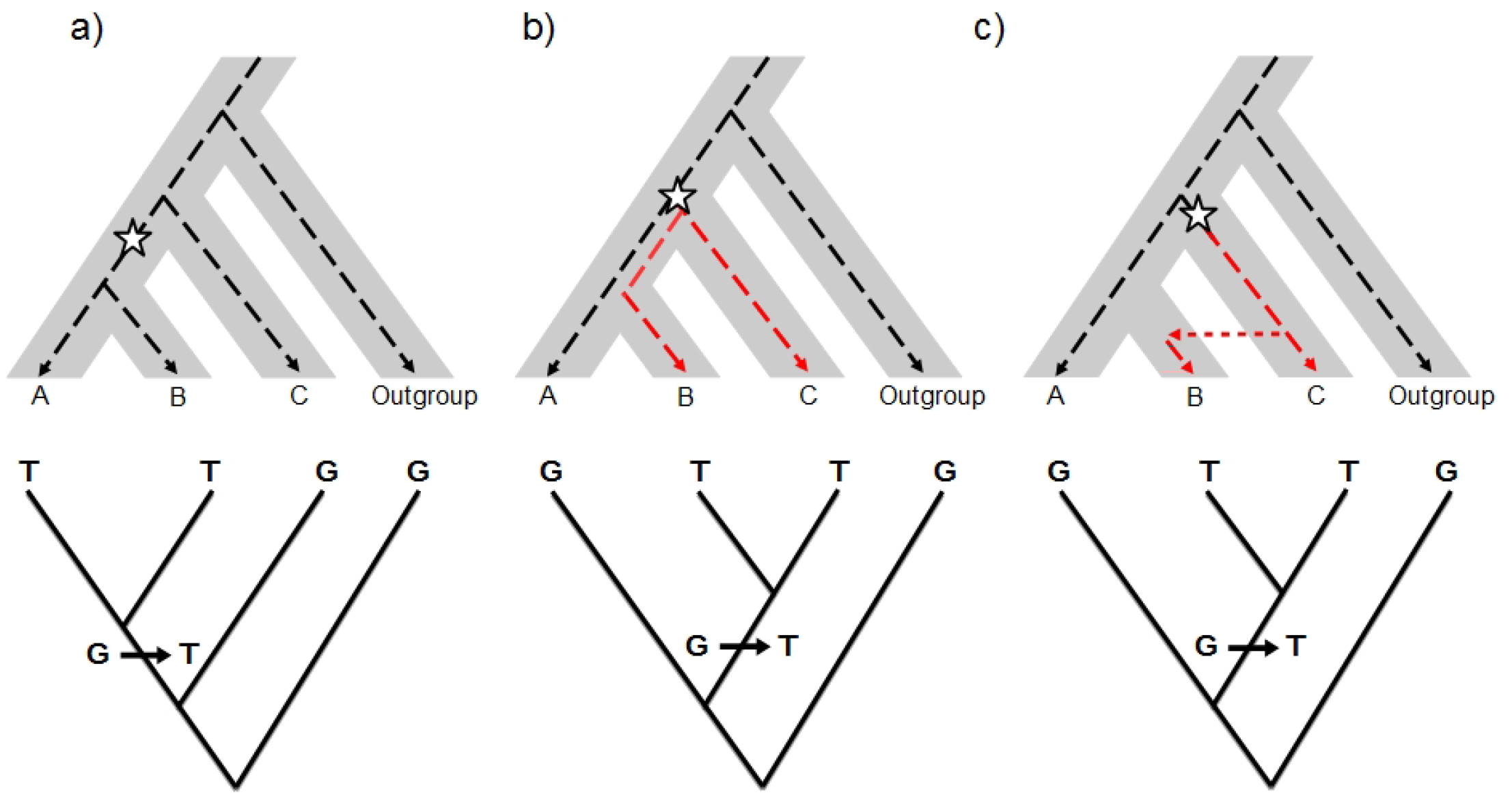
Hybridization and incomplete lineage sorting (ILS) revealed by molecular phylogenetics. Top: evolutionary process. The species (or population) tree is represented by the grey area. The dotted line represents a single gene genealogy. Bottom: Coalescent tree inferred for the gene. (a) Congruent gene genealogy with species/population tree; (b) ILS: ancestral polymorphism is maintained before the divergence between A and B, so that B shares the allele T with C and not with A; (c) Introgression: B receives the allele T from C by gene flow. In the case of ILS and introgression, the gene genealogy is incongruent with the species/population tree.

Coalescent samplers have been widely used to test for gene flow versus ILS using maximum likelihood or Bayesian models (Pinho and Hey, 2010). However, they are not straightforwardly applied to all study systems, because they require a strong computation effort and are not easy to transpose to a genome-wide scale. An alternative, simpler strategy takes advantage of the expectations associated with phylogenetic relationships between individuals or populations (Figure 3). Given a genealogical tree describing the history of divergence between taxa or populations, a precise amount of shared variation between branches is expected because of drift and ILS. A significant excess of shared variation instead may be indicative of gene flow (Kulathinal et al., 2009; Patterson et al., 2012; Peter, 2016). A number of statistics have been developed to test for the excess of shared polymorphism. The most used are the D-statistic (or ABBA-BABA test, Durand et al., 2011; Green et al., 2010) and the *f3* and *f4* statistics (globally referred to as *f*-statistics, Reich et al., 2009, 2012). These statistics were initially applied to human populations and have proven to be useful in other study systems, e.g. to detect the introgression of adaptation to serpentine soils in *Arabidopsis* species (Arnold et al., 2016). In general, the power of these tests to detect admixed genomes or populations is greater when applied to genome-wide data (see Patterson et al., 2012 for a review; Peter, 2016), but most recent statistics can be applied to small genomic regions, e.g. *f*_D_ (Martin et al., 2015; Racimo et al., 2017).

Other approaches take advantage of haplotype characteristics to distinguish between introgression and ILS. As recombination breaks apart haplotypes over generations, introgressed haplotypes should be longer than haplotypes due to ILS and should exhibit higher levels of linkage disequilibrium (see figure 1 from Racimo et al., 2015). If admixture occurred recently compared to the divergence between populations, these features can be exploited to detect introgressed tracts. A test of significance can be associated by performing coalescent simulations of specific demographic scenarios (setting values of divergence times, recombination rates, population structure or selection adapted to the case in hand) to obtain the expectations for haplotype length statistics in the absence of gene flow. Haplotype length analyses led to the identification of candidate introgressed tracts and estimation of the age of the last introgression event in humans (Racimo et al., 2015) and dogs (Miao et al., 2016). A recently developed statistic, S*, uses linkage disequilibrium information to detect introgressed haplotypes when no information about the donor is available. S* is designed to identify divergent haplotypes whose variants are in strong linkage disequilibrium and are not found in a non-admixed reference population. S* increases as the number of linked SNPs and the distance between them increases (Vernot et al., 2016). This statistic helped to reveal the introgressed origin of the EPAS1 gene in Tibetans, before the identification of the Denisovan donor (Huerta-Sánchez et al., 2014).

### 3.2 Detection of selection

To prove adaptive introgression, the action of selection has to be demonstrated on the introgressed variant. A number of reviews address methods and tools for detecting selection with molecular data (e.g. Bank et al., 2014; Pavlidis and Alachiotis, 2017). In practice, most of the available approaches are more sensitive to signatures of strong positive selection (i.e. selective sweeps, Smith and Haigh, 1974). For regions under strong positive selection, expectations are lower diversity, higher linkage disequilibrium and specific distortions of the allele frequency spectrum compared to the genome-wide patterns.

In within-population analyses, local patterns of lower genome diversity (Figure 2b) and shifts of the allele frequency spectrum toward an excess of low frequency alleles are often informative for detecting positive selection. For instance, polymorphism summary statistics, such as *π* (nucleotide expected heterozygosity) and Tajima’s D, have helped to discover and characterize introgressed loci involved in serpentine adaptation of *Arabidopsis arenosa* (Arnold et al., 2016) and in the pesticide resistance of mosquitoes (Norris et al., 2015) and mice (Song et al., 2011). Advanced methods for genomic scans of positive selection are the Composite Likelihood Ratio test approaches (reviewed in Pavlidis and Alachiotis, 2017). These tests compare the probability of the observed local site frequency spectrum under a model of selection with the probability of observing the data under the standard neutral model. The neutral expectations can be inferred by genome-wide observed patterns or by specific simulated demographic scenarios (e.g. Liu et al., 2015; Quach et al., 2016; Staubach et al., 2012).

Haplotypic information is also extremely useful for identifying almost fixed or very recently fixed selective sweeps. The frequency of the introgressed haplotype in the recipient population can serve for identifying selection. This interpretation is based on the assumption that introgressed regions under selection should be at higher frequencies in the population relatively to the rest of the genome (e.g. Vernot et al., 2016). The extent of linkage disequilibrium generated on the sides of a beneficial mutation (or the haplotype size) is another signature captured by a number of tests for selection (Crisci et al., 2012). The BADH2 gene, responsible for the much-appreciated characteristic fragrance of some Asian rice varieties, provides a nice example of adaptive introgression detected by haplotype analysis. This gene only shows strong signatures of selection in fragrant accessions, as revealed by a dramatic reduction in diversity (*π*) and a large block of linkage disequilibrium in regions flanking the functional mutation. The selected fragrant allele is likely to have originated after domestication in the genetic background of the *japonica* varietal group and to have been transferred to the *indica* variety by introgression (Kovach et al., 2009).

Extreme differentiation between populations in specific genomic regions can also be interpreted as a signature of selection subtending local adaptation. For introgressed alleles adaptive in the recipient population, higher differentiation can be expected between the recipient and another non-admixed population (e.g. Ai et al., 2015). In addition, recipient-donor differentiation will be lower for introgressed regions compared to the rest of the genome (Figure 2b). Thus, comparisons of pairwise differentiation values between different populations (i.e. donor, recipient and “reference” non-admixed population) may help to disentangle instances of adaptive introgression (e.g. Arnold et al., 2016; Enciso-Romero et al., 2017; Racimo et al., 2017). A number of differentiation/divergence statistics with different properties are available (e.g. Cruickshank and Hahn, 2014). Among them, estimators of *F_ST_* (Wright, 1931) are the most commonly used for detecting selection (e.g. Arnold et al., 2016; Gagnaire et al., 2009; Gittelman et al., 2016).

It should be noted, however, that inferring separately introgression and selection might not be the best approach to detect adaptive introgression, as expected genetic patterns are not necessarily the same. For example, we do not necessarily expect a reduction of diversity in an introgressed region under positive selection. In fact, it has been shown by simulations that admixture can increase diversity blowing the diversity loss due to selection (Racimo et al., 2017). Recent investigations into the joint dynamics of introgression and positive selection have opened promising avenues for the analysis of genetic data in quest of adaptive introgression instances (Racimo et al., 2017). These authors proposed new statistics informative to identify candidates to adaptive introgression and based to the number and frequency of alleles shared by the donor and the recipient populations (but absent or nearly absent in non-introgressed reference populations). Such “unique shared alleles” should be numerous and at high frequency in genomic regions interested by adaptive introgression (Figure 3c). The proposed statistics resuming these patterns, Q95 and U, have proven successful to retrieve several known regions of archaic adaptive introgression from Neanderthals and Denisovans in modern human genome (Racimo et al., 2017). However, these statistics are not straightforwardly applicable to any study system. Specific demographic simulations are necessary to assess their expected value in absence of adaptive introgression.

Different types of selection other than selective sweeps may generate genetic patterns that are more difficult to distinguish from non-selective processes with the approaches described above. Balancing selection on an introgressed locus, for instance, might go undetected because it would maintain several variants at intermediate frequency within the recipient population. Such a pattern can be interpreted as the result of migration-drift equilibrium, unless a direct link has been established between the locus and a phenotypic trait. Examples of adaptive introgression driven by balancing selection are the incompatibility locus in *Arabidopsis* (Castric et al., 2008), skin colour change in wolves (Anderson et al., 2009; von Holdt et al., 2016) and the HLA locus in humans (Abi-Rached et al., 2011). Soft-sweeps, fixation of a beneficial allele starting from multiple copies of it in the population (Hermisson and Pennings, 2005), could also interest introgressed regions. Typically, when the migration rate is high, the same beneficial allele can enter the recipient population associated with different genetic backgrounds. This kind of positive selection signature is difficult to detect because diversity and site frequency spectrum patterns do not change dramatically as in hard selective sweeps (Hermisson and Pennings, 2017).

It should be noted also that inferences of selection based on molecular data only give indirect evidence of the adaptive value of introgressions, particularly when they target genomic regions with an unknown contribution to fitness-related traits (e.g. Gagnaire et al., 2009). However, detecting selection in genic regions linked to specific functions or phenotypes (shown by phenotype-genotype association analysis for instance) greatly helps the interpretation in terms of adaptation (e.g. Hufford et al., 2013; Racimo et al., 2015; Rochus et al., 2017). Ultimately, one direct validation of the adaptive role of introgression is to demonstrate the fitness advantage of the introgressed allele or trait for the recipient population (e.g. Martin et al., 2006; Whitney et al., 2006, 2010, 2015; Figure 4). However, field studies involving phenotypic exploration can be time-consuming and difficult to implement for most species.

**Figure 4.**
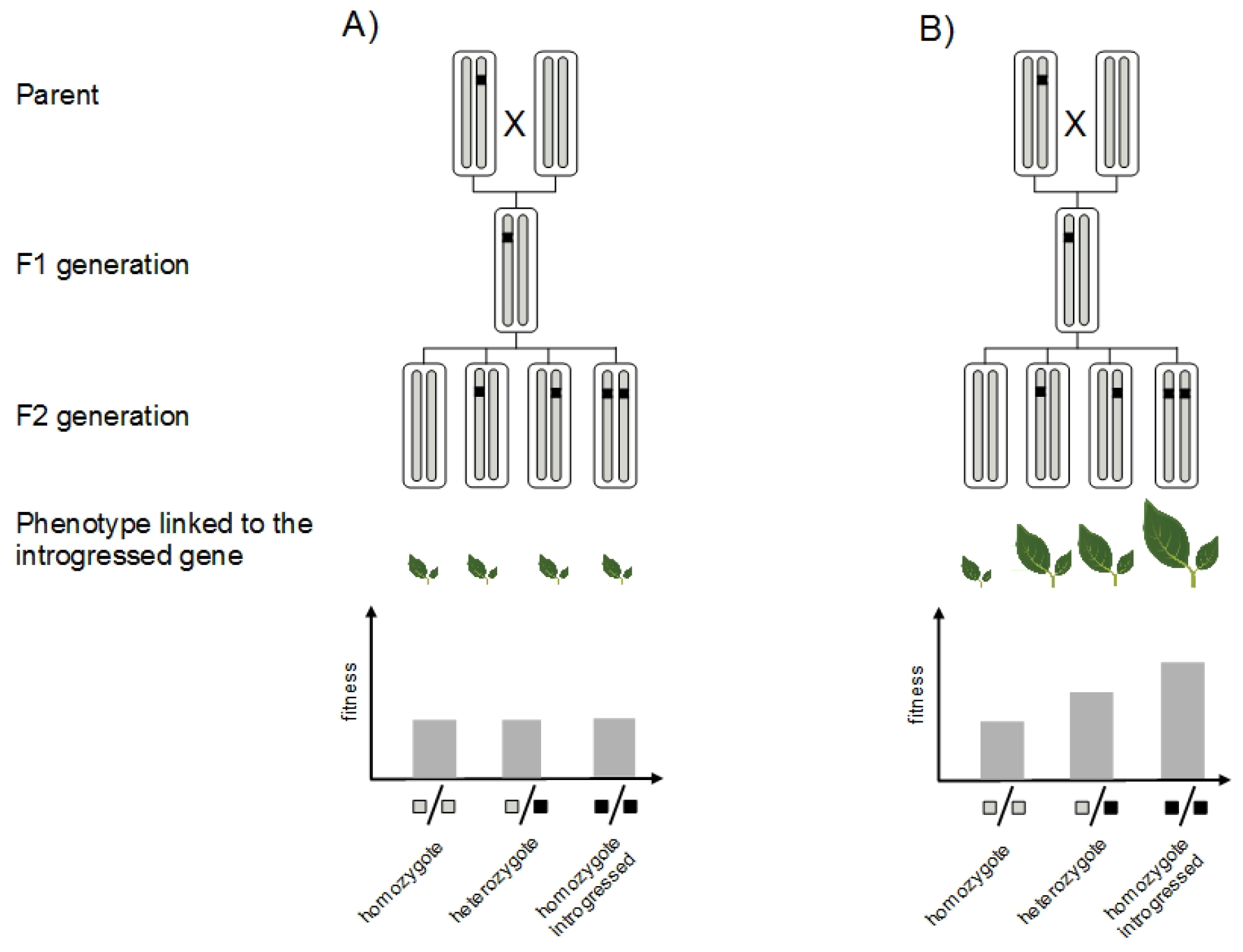
Direct measure of adaptive introgression. Direct evidence of the adaptive value of the introgressed fragment (black segment) consists in showing that it confers greater fitness to the recipient genome. This can be achieved by experimental crosses: introgression without positive selection on the introgressed allele (A) *vs* adaptive introgression (B).

## 4. Gene flow in crops: challenges and opportunities

The domestication process often involves dispersion across long distances and different environmental conditions, combined with intense selection. Under such a process, multiple opportunities arise for gene flow with locally adapted cultivated or wild forms (Allaby et al., 2008; Meyer and Purugganan, 2013). In addition, among domestication characters, genetic divergence between domesticated and wild relatives is often low and the reproductive barriers narrow (e.g. Dempewolf et al., 2012). These characteristics make domesticated species likely carriers of adaptive introgression.

Thanks to the analysis of genome-wide data, the importance of gene flow in shaping today’s diversity of domesticated species is becoming evident, particularly in crops. The evolution of two major cereals, rice and barley, fits a domestication history with frequent gene flow events. In rice, complex introgressive hybridizations have shuffled the genome of Asian rice, leading to the current main groups *O. sativa japonica* and *O. s. indica*. Surveys indicate that *O. s. indica* had acquired major domestication alleles through gene flow from *O. s. japonica* into the wild progenitor *O. rufipogon*, or into putative proto-*indica* populations (Choi et al., 2017). White pigmentation, aromatic fragrance and glutinous starch are some of the phenotypic traits involved in allele transfers driven by strong directional selection (Huang et al., 2012; Kovach et al., 2009; Olsen and Purugganan, 2002). In barley, the genome of domesticated forms appears to be a mosaic of fragments originating from different cultivated and wild populations across the Fertile Crescent (Pankin and von Korff, 2017; Poets et al., 2015). Introgression from local wild relatives has also been shown in grapes (Myles et al. 2011), apples (Cornille et al., 2012) and olives (Diez et al., 2015). Similarly, gene flow events have occurred in domesticated animals. For instance, Rochus et al. (2017) analyzed the diversity of French sheep breeds to find that one mutation in a genomic region related to milk production and growth originated in a southern breed and was introgressed and selected for in northern breeds.

While adaptive introgression of agronomic traits between different breeds or cultivars may occur relatively easily under similar human selective pressures, the transfer of beneficial traits from wild relatives seems more difficult, since selection in natural environments usually goes in the opposite direction from selection in agricultural environments. Crop-wild admixed plants (weeds) may tend to be phenotypically intermediate and thus to have unwanted wild-type traits (e.g. asynchrony of phenology changes, multiple branching, seed shattering, noxious or unpalatable compounds). To understand how wild alleles can enter the crop gene pool by spontaneous gene flow, Jarvis and Hodgkin (1999) point to the necessity of understanding how farmers’ taxonomy and practices (i.e. seed management, weeding, etc.) are applied to the new phenotypic variations resulting from hybridization. Barnaud et al. (2009) investigated criteria used by farmers to characterize morphotypes of domesticated and weedy sorghum. While strong counter-selection was performed against weedy types, they found that progenies of weedy types could be misidentified as cultivated forms, thus favoring wild-to-crop introgression. For pearl millet, it has been suggested that incomplete weeding and singling allow hybridization and introgression to occur freely and extensively (Couturon et al., 1997; Robert et al., 2003, Mariac et al. 2006), favoring the maintenance of wild genetic material in the cultivated gene pool. In addition, weedy types can play an important role for food security. In many cases, weedy types are early maturing plants and they are used under harsh conditions or between main harvests. In Sudan, farmers recognize a crop-wild hybrid of sorghum, which is allowed to grow and is selectively harvested in bad years (Ejeta and Grenier, 2005). Hybrids can be harvested during periods of scarcity in the case of pearl millet too (Couturon et al., 2003; Mariac et al., 2006) or in common bean (de la Cruz et al., 2005; Zizumbo-Villarreal et al., 2005). Some studies have also documented farmers practicing conscious directional selection towards evolution and changes of cultivated phenotypes by using the diversity available in the wild relatives. For instance, in Benin, farmers voluntarily grow wild and hybrid yams (*Dioscorea* spp.) in their fields to increase diversity (Scarcelli et al., 2006).

All in all, this demonstrates how farmer practices can maintain and, in some cases, actively favor wild-to-crop introgression. This can be particularly important for crop adaptation, since the trade-off of strong human selection for certain traits has been the loss of diversity for other important adaptive traits (e.g. Zheng et al., 2008). Notably, traits involved in climate and soil adaptation, or resistance to pests and diseases, display much greater diversity in wild species than in domesticated species (Dempewolf et al., 2017; Guarino and Lobell, 2011; Hajjar and Hodgkin, 2007). Therefore, favoring introgression from wild relatives may be an indirect way for farmers to introduce lost or new adaptations. A compelling example of the potential adaptive outcome of wild-to-crop introgression is the adaptation to altitude acquired by highland maize landraces. Maize was domesticated from low altitude wild populations of teosinte (*Zea mais ssp.parviglumis*) and colonized high altitude environments (Matsuoka et al., 2002) where a different wild relative is found (Z. m. *mexicana*). Hufford et al. (2013) performed genomic scans on Mexican sympatric populations of maize and *mexicana* and found nine genomic regions of introgression of *mexicana* into maize landraces. These regions, related to adaptive traits, such as the quantity of leaf macrohairs and pigmentation intensity, could have helped maize to adapt to high altitude (Hufford et al., 2013). Recent study suggested that wild-to-crop gene flow significantly genetic diversity and possibly lead to introgressions of local adaptation in pearl millet, a major staple African crops (Burgarella et al., in press).

The transferal of wild traits into a cultivated genetic background through classic breeding programs has so far been limited to a narrow range of traits (Warschefsky et al., 2014), e.g. disease resistance in cassava (Bredeson et al., 2016) and tomatoes (Lin et al., 2014). One main difficulty for conventional breeding in introgressing wild genes into cultivated gene pools is that wild diversity is mostly unexplored and desirable characters have still to be identified. Furthermore, wild species have many traits associated with poor agronomic performance (e.g. low yields, seed shattering, small seed or fruit size). Thus, several backcrosses are necessary to dilute the unwanted diversity associated with wild introgression (Dempewolf et al., 2017) or to produce specifically design lines in which small regions of the wild relative are introduced (e.g. Fonceka et al., 2012). We argue that screening the wild introgression already existing in the cultivated gene pool thanks to natural gene flow may be an effective strategy to uncover wild diversity relevant for crop adaptation to environmental changes and to inform new breeding directions. Therefore, research efforts should be devoted to quantifying and characterizing the extent of such spontaneous wild-to-crop introgression. Two main challenges are associated with this approach. First one will need to have access to wild “pure” genetic resources, which highlight the crucial need of wild relative conservation programs. Second, identification of adaptive introgression in crops might be more challenging because of the low genetic divergence usually observed between crop species and their wild relatives (as exposed in paragraph 3). However, interest in the study of adaptive introgression, notably in humans, means the field is moving fast towards new analytical approaches designed to identify specific features in genomes (e.g. Racimo et al., 2017). Of course, the validation of adaptive introgression detected via molecular data should be validated with classic experiments (Figure 4) to measure the strength of selection in the field and to assess the biological function of the introgressed alleles (Suarez-Gonzalez et al., 2018; Whitney et al., 2006). Given their adaptation to human-controlled environments, this step seems easier to accomplish in most crops than in wild species.

Regarding the introduction or re-introduction of adaptive variation, it might be wondered to what extent introgression from wild relatives can affect the whole crop genome. Recent studies have suggested that introgression can be favored at genome-wide level when it reduces the genetic load of the recipient species (Sankararaman et al., 2014; Wang et al., 2017). Genetic load refers to the genome-wide accumulation of weakly deleterious alleles that reduces its fitness (Crow, 1958). Given the repeated selection rounds associated with the domestication process, crop species experience a reduction in the effective population size and in effective recombination, which in turn reduces the efficacy of purifying selection in removing deleterious alleles and increases the effect of hitchhiking selection (i.e. deleterious variants increase in frequency because they are linked to selected beneficial alleles). Inbreeding, which is commonly practiced to fix traits of interest, also slightly contributes to fixing deleterious alleles. In crops, a reduction in fitness is expected compared to the wild progenitor, the so-called ‘cost of domestication’ (Lu et al., 2006). A greater genetic load than in the wild counterpart was observed in several domesticated species such as rice (Lu et al., 2006), maize (Wang et al., 2017), sunflower (Renaut and Rieseberg, 2015), dogs (Marsden et al., 2016) and horses (Schubert et al., 2014). Since wild species are expected to have a lower genetic load than cultivated species, spontaneous introgression from wild species could be favored, because it alleviates the domestication cost, even in the absence of strong directional selection on the introgressed alleles. Recent findings in maize support this expectation, as negative correlations were observed between wild introgression and genetic load (Wang et al. 2017).

Despite the potential benefits of wild introgression for crops (i.e. acquisition of specific adaptations and reduction of genetic load), genomic heterogeneity is expected in terms of permeability to gene flow. In particular, regions involved in major domestication characters are expected to be under strong selection, thereby acting as barriers to gene flow (so-called “islands of domestication”, Frantz et al., 2015). Thus, the probability of introgression along the crop genome largely depends on the number and distribution of domestication loci. Loci responsible for domestication traits have been identified in a number of crops (Doebley et al., 2006; Gross and Olsen, 2010; Meyer et al., 2012), but knowledge is far from complete. Up to now, research on the genetic architecture of domestication traits indicates that domestication loci are limited to a few genomic regions in most studied species (Burger et al., 2008; Glémin and Bataillon, 2009) and may not be a major obstacle to introgression in the rest of the genome. The efficiency of counter-selection would thus depend on the genetic distance between the introgressed fragment and the domestication genes, which is determined by the extent of local linkage disequilibrium. According to this expectation, Hufford et al. (2013) identified cold spots and hotspots of wild introgression in the maize genome. Interestingly, cold spots were significantly enriched in domestication genes (Hufford et al., 2013).

## 5. Conclusion and prospects

Today’s access to both phenotypic and genomic information provides the opportunity to further investigate the role and mechanism of adaptive introgression in crops. From an applied point of view, it can be a fair source of adaptation to be exploited in breeding programs. Along human migrations, crops had the opportunity to exchange with multiple wild populations. These exchanges might have resulted in the introgression of local adaptations that have already passed the reproductive and the “agronomic” barriers. Identifying such local adaptive introgression, combined with complementary tools (e.g. climatic modeling), could be an efficient way of adapting crops to the predicted new environmental conditions. We therefore think there is a need to emphasize the importance of conserving wild genetic resources and jointly investigating wild and crops relatives. This research will also allow advances on key questions with broader prospects, such as introgression rates and genome permeability (Hufford et al., 2013; Scascitelli et al., 2010). Genomic scan approaches could be complemented by the development of recombinant inbred line (RILs) wild-crop hybrid populations (e.g. Fonceka et al., 2012; Nice et al., 2016). Such pre-breeding material can be exchanged and tested in different environments, which would also help to answer the other side of the coin, the risk of transgene escapes in the case of crop-to-wild gene flow.

## Acknowledgements

We thank Peter Biggins for English revision and Miguel Navascués for useful discussions on methodological approaches. This project was supported by Agropolis Fondation under the reference ID 1403-057 through the « *Investissements d’avenir* » programme (Labex Agro:ANR-10-LABX-0001-01) under the frame of I-SITE MUSE (ANR-16-IDEX-0006 and the the NERC/DFID Future Climate For Africa programme under the AMMA-2050 project, grant number NE/M019934/1.

## Author contribution statement

CB, AB and CB-S wrote the first draft, all authors made a substantial, direct and intellectual contribution to this work.

